# Metabolic Recovery and Compensatory Shell Growth of Juvenile Pacific Geoduck *Panopea Generosa* Following Short-Term Exposure to Acidified Seawater

**DOI:** 10.1101/689745

**Authors:** Samuel J. Gurr, Brent Vadopalas, Steven B. Roberts, Hollie M. Putnam

## Abstract

While acute stressors can be detrimental, environmental stress conditioning can improve performance. To test the hypothesis that physiological status is altered by stress conditioning, we subjected juvenile Pacific geoduck, *Panopea generosa*, to repeated exposures of elevated *p*CO_2_ in a commercial hatchery setting followed by a period in ambient common garden. Respiration rate and shell length were measured for juvenile geoduck periodically throughout short-term repeated reciprocal exposure periods in ambient (~550 µatm) or elevated (~2400 µatm) *p*CO_2_ treatments and in common, ambient conditions, five months after exposure. Short-term exposure periods comprised an initial 10-day exposure followed by 14 days in ambient before a secondary 6-day reciprocal exposure. The initial exposure to elevated *p*CO_2_ significantly reduced respiration rate by 25% relative to ambient conditions, but no effect on shell growth was detected. Following 14 days in common garden, ambient conditions, reciprocal exposure to elevated or ambient *p*CO_2_ did not alter juvenile respiration rates, indicating ability for metabolic recovery under subsequent conditions. Shell growth was negatively affected during the reciprocal treatment in both exposure histories, however clams exposed to the initial elevated *p*CO_2_ showed compensatory growth with 5.8% greater shell length (on average between the two secondary exposures) after five months in ambient conditions. Additionally, clams exposed to the secondary elevated *p*CO_2_ showed 52.4% increase in respiration rate after five months in ambient conditions. Early exposure to low pH appears to trigger carry-over effects suggesting bioenergetic re-allocation facilitates growth compensation. Life stage-specific exposures to stress can determine when it may be especially detrimental, or advantageous, to apply stress conditioning for commercial production of this long-lived burrowing clam.

**Lay summary:** Commercial shellfish hatcheries provide essential food security, but often production can be hampered by sensitivity of shellfish at early life stages. Repeated short-term exposures can increase tolerance and performance of the geoduck clam with implications for sustainable aquaculture.

## 1. Introduction

Sustainable food production minimizes overexploitation of wild populations and degradation of ecological health (Campbell *et al.*, 1998; Shumway *et al.*, 2003; Orensanz *et al.*, 2004; Zhang and Hand, 2006). Shellfish aquaculture has expanded worldwide in recent decades to satisfy international trade (FAO 2018). However, early larval and juvenile rearing pose a production bottleneck. For example, early life histories are highly sensitive to biotic (e.g. harmful algae, pathogens; Prado *et al.*, 2005; Rojas *et al.*, 2015) and abiotic stressors (e.g. pH, salinity, thermal, and hypoxic stress; Baker and Mann 1992; Przeslawski et al. 2015; Kroeker *et al.*, 2010; Gimenez *et al.*, 2018). These stressors are known to intensify in coastal marine systems (Cloern, 2001; Diaz and Rosenberg, 2001; Cai *et al.*, 2011; Wallace *et al.*, 2014) causing mass mortality for early-stage bivalves in wild or hatchery settings (Elston *et al.*, 2008; Barton *et al.*, 2015). Local and global anthropogenic stressors such as CO_2_-induced changes in pH and carbonate mineral saturation states can reduce performance and normal shell development (White *et al.*, 2013; Waldbusser *et al.*, 2015; Kapsenberg *et al.*, 2018).

Ocean acidification, or the decrease of oceanic pH due to elevated atmospheric partial pressures (µatm *p*CO_2_), poses a threat to aquaculture (Barton *et al.*, 2012; Froehlich *et al.*, 2018; Mangi *et al.*, 2018). Elevated *p*CO_2_ and aragonite undersaturation (Ω_aragonite_ < 1) generally have detrimental consequences for aerobic performance (Pörtner *et al.*, 2004; Portner and Farrell, 2008) and shell biomineralization in marine calcifiers (Shirayama, 2005; Talmage and Gobler, 2010; Waldbusser *et al.*, 2010, 2015; Gazeau *et al.*, 2013). Responses to acidification can be species (Ries *et al.*, 2009) and population specific (Lemasson *et al.*, 2018), but it is widely established to be impactful during early life stages for bivalves (Dupont and Thorndyke, 2009; Gazeau *et al.*, 2010; Kroeker *et al.*, 2010; Gimenez *et al.*, 2018). Experimental research is commonly focused on species with short generational times, (Parker *et al.*, 2011, 2015; Lohbeck *et al.*, 2012) limiting evidence for effects of acidification on long-lived mollusks important for food and economic security (Melzner *et al.*, 2009).

The Pacific geoduck *Panopea generosa* is a large and long-lived infaunal clam of cultural and ecological importance (Dethier, 2006) with an increasing presence in sustainable shellfish industry (Cubillo *et al.*, 2018). Geoduck production in Washington (USA) provides ~90% of global supply (Shamshak and King, 2015) and alone constitutes 27% of the overall shellfish revenue in the state valued at >$24 million yr^−1^ and >$14 pound^−1^ as of 2015 (Washington Sea Grant, 2015). Geoduck are known to live in dynamic CO_2_-enriched low pH waters such as Hood Canal in Puget Sound, WA where conditions in summer can reach Ω_aragonite_ 0.4 and pH 7.4 (Feely *et al.*, 2010). Although *P. generosa* may be adapted and able to acclimatize to local stressors (Putnam *et al.*, 2017; Spencer *et al.*, 2018), acidification has caused massive losses of larval bivalves in hatcheries (Barton *et al.*, 2015), identifying a critical need for assessment of physiological stress tolerance during early life stages.

Evidence of acclimatory mechanisms in response to acidification (Goncalves *et al.*, 2018) and enhanced performance within and across generations (Parker *et al.*, 2011, 2015; Putnam and Gates, 2015; Ross *et al.*, 2016; Thomsen *et al.*, 2017; Zhao *et al.*, 2017) support conditioning as a viable strategy to mitigate the negative effects of stress exposure and enhance organismal performance under high *p*CO_2_ (Parker *et al.*, 2011; Dupont *et al.*, 2012; Suckling *et al.*, 2015; Foo and Byrne, 2016). Hormesis is a biphasic low-dose-stimulatory response, as identified in toxicological studies (Calabrese, 2008), and suggests beneficial carry-over effects of moderate stress exposure (Calabrese *et al.*, 2007; Costantini *et al.*, 2010; Costantini, 2014; Putnam *et al.*, 2018). Conditioning-hormesis can explain patterns of intra- and transgenerational plasticity for organisms under environmental change (Calabrese and Mattson, 2011; Costantini *et al.*, 2012; López-Martínez and Hahn, 2012; Putnam *et al.*, 2018; Visser *et al.*, 2018), but is understudied for stress resilience in bivalves likely due to generally negative physiological implications of acidification (Gazeau *et al.*, 2013). In one example of early-life stage conditioning in bivalves, Putnam et al. (2017) found *P. generosa* exhibit compensatory shell growth after an acute exposure under elevated *p*CO_2_. This finding suggests acute exposures may present a strategy for stress-hardening and enhancement of sustainable geoduck production. We therefore tested the hypothesis that repeated stress exposure under elevated *p*CO_2_ can enhance intragenerational performance for Pacific geoduck. To this end, we measured the respiration rate and shell growth of juvenile geoduck in a commercial hatchery under repeated acute periods (~6-10 days) of elevated *p*CO_2_ and aragonite undersaturation, and the longer term (~5 months) carry-over effects.

## 2. Methods

### 2.1. Exposure of juveniles

Juvenile geoduck (n = 640; mean ± SEM initial size, 5.08 ± 0.66 mm shell length [measured parallel to hinge]) were reared in trays (Heath/Tecna water tray) with rinsed sediment for ~16 weeks (pediveliger to juvenile stage) by Jamestown Point Whitney Shellfish Hatchery before allocated into eight trays for the experiment (Fig. 1; n = 80 clams per tray). During typical hatchery practice, geoduck are reared from ‘settlers’ (pediveliger stage; 30 days old) to ‘seed’ (juvenile stage; 4-6 months old) in either downwellers or stacked trays; juveniles are then planted *in-situ* to grow for several years until market size. Following aquaculture practice, trays were filled with 5 mm depth of rinsed sand (35-45 µm grain size) that allowed juvenile geoduck to burrow and siphons could clearly be seen extended above the sediment throughout the experiments. To enable measurements of metabolic activity and shell growth, 30 geoduck were placed in an open circular dish (6.5 cm diameter and 3 cm height) with equal mesh size and sand depth submerged in each tray, the remaining 50 geoduck in each tray burrowed in the surrounding sediment. Seawater at the Jamestown Point Whitney Shellfish Hatchery (Brinnon, WA, USA) was pumped from offshore (100 m) in Quilcene Bay (WA, USA), bag-filtered (5 µm), and UV sterilized before fed to 250-L conical tanks at rate of 1 L min^−1^. Four conical tanks were used as replicates for two treatments: elevated *p*CO_2_ level of ~2300-2500 µatm and ~7.3 pH (total scale); and ambient hatchery conditions of ~500-600 µatm and ~7.8-7.9 pH (total scale). The elevated *p*CO_2_ level was set with a pH-stat system (Neptune Apex Controller System; Putnam *et al.*, 2016) and gas solenoid valves for a target pH of 7.2 (NBS scale) and pH and temperature (°C) were measured every 10 seconds in conicals (Neptune Systems; accuracy: ± 0.01 pH units and ± 0.1°C, resolution: ± 0.1 pH units and ± 0.1°C). These treatments were delivered to replicate exposure trays, which were gravity fed seawater from conicals (Fig. 1; n = 4 per treatment). The experiment began with an initial exposure period of 10 days under elevated *p*CO_2_ (2345 µatm) and ambient treatments (608 µatm; Table 1). Preliminary exposure was followed by 14 days in ambient common garden (557 ± 17 µatm; pH_t.s._ 7.9 ± 0.01; Ω_aragonite_ 1.46 ± 0.04, mean ± SEM) before secondary exposure for 6 days to reciprocal treatments of elevated *p*CO_2_ (2552 µatm) and ambient treatments (506 µatm; Table 2). For the secondary exposure period, one tray was crossed to the opposite treatment to address both repeated and reciprocal exposure (n = 2 trays per initial×secondary *p*CO_2_ treatment; Fig. 1). Following this the juveniles were exposed to ambient conditions for 157 days within the replicate trays.

**Table 1.**
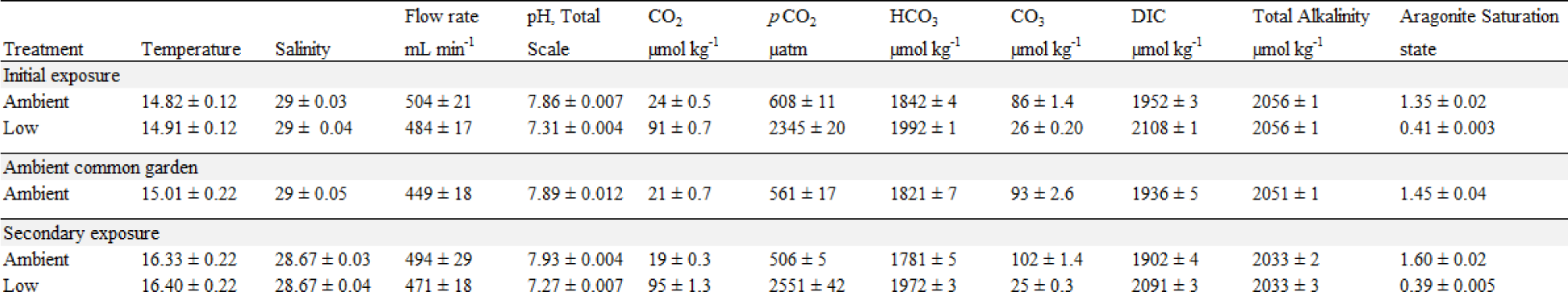
pH, salinity, and temperature measured with handheld probes and total alkalinity measured daily with 60 mL from each heath tray via Gran titration (n = 4 per treatment) during initial (10-d) and secondary (6-d) exposure trials. Seawater carbonate chemistry (CO_2_, *p*CO_2_, HCO_3_^−^, CO_3_^2-^, DIC, aragonite saturation state) was calculated with the seacarb R package (Gattuso *et al.*, 2018).

**Table 2.**
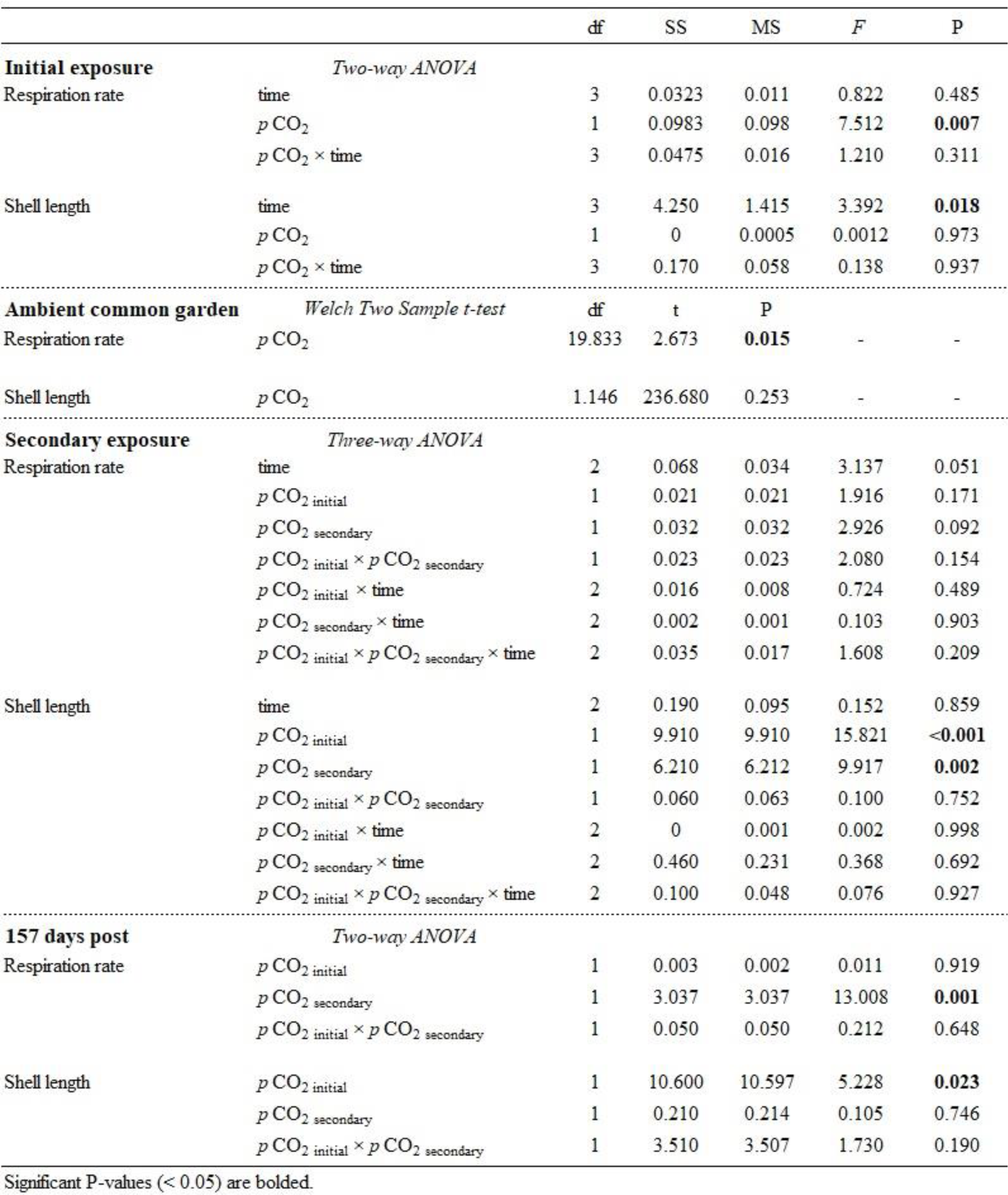
Two-way and three-way ANOVA tests for metabolic rate and shell length during initial and secondary exposures, respectively. A Welch’s t-test was used on day zero of secondary exposure to test for differences in mean respiration rate and shell length from initial treatments and a two-way ANOVA tested for treatment effects after 157 days. Significant effects are bolded for P < 0.05.

**Figure 1.**
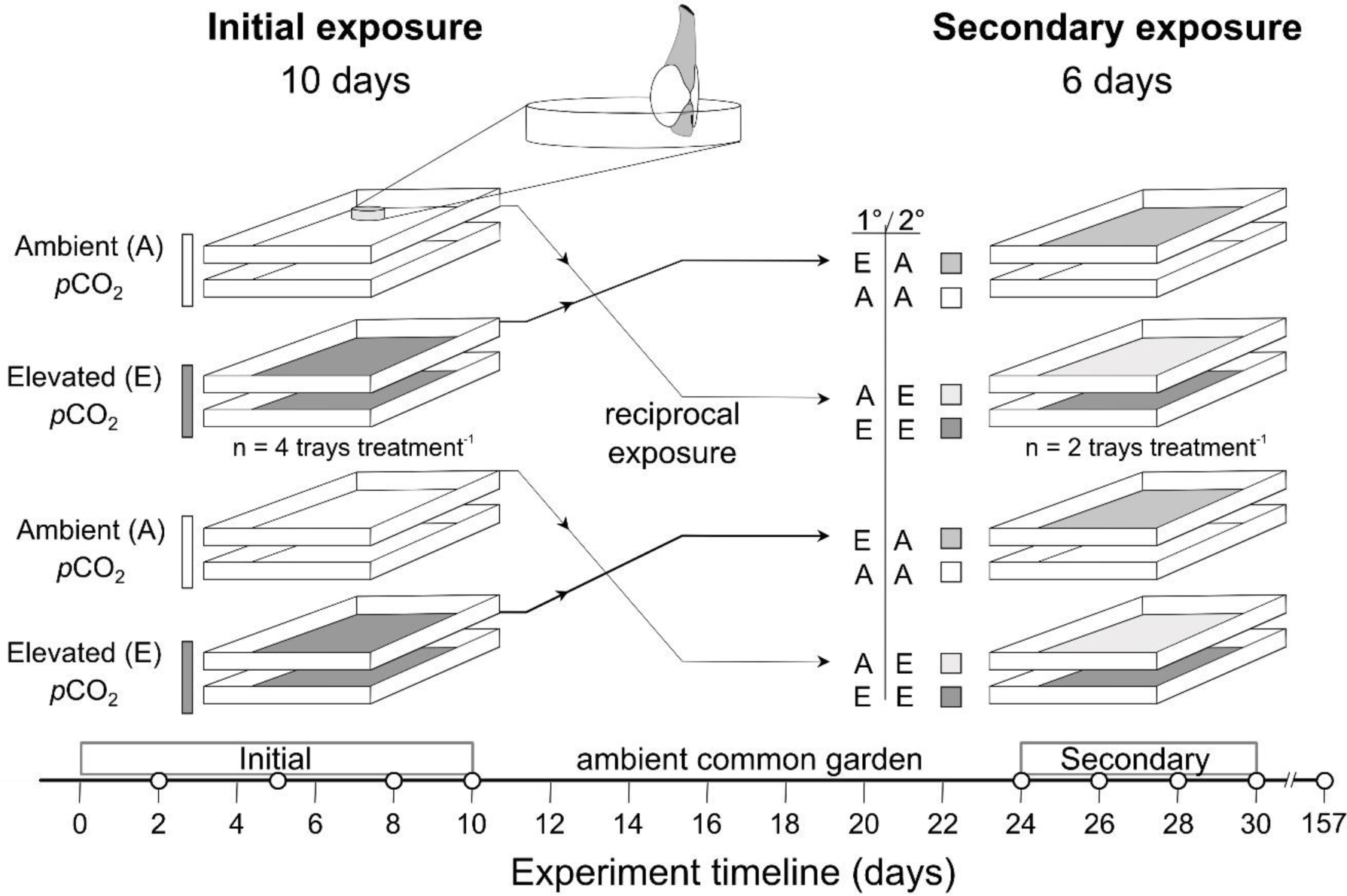
Schematic of the repeated exposure experimental design for two exposure trials, initial (10-day) and secondary (6-day), in ambient and elevated *p*CO_2_ treatments. Timeline displays respiration and growth measurements as solid white circles.

Juvenile geoduck were fed semi-continuously with a mixed algae diet (30% *Isocrysis galbana*, 30% *Pavlova lutheri*, and 40% *Tetraselmis suecica*) throughout the 30-d experiment with a programmable dosing pump (Jebao DP-4 auto dosing pump). Large algae batch cultures were counted daily via bright-field image-based analysis (Nexcelom T4 Cellometer; Gurr *et al.*, 2018) to calculate a daily ration of 5×10^7^ live algae cells d^−1^ individual^−1^. Diet was calculated with an equation in Utting & Spencer (1991) catered for 5-mm clams:

V = (S × 0.4) ÷ (7 × W × C); this equation accounts for a feed ration of 0.4 mg dried algae mg live animal weight^−1^ week^−1^, the live animal weight (mg) of spat (S; estimated from regression of shell length and weight of Manilla clams in Utting and Spencer 1991), weight (mg) of one million algal cells (W), and cell concentration of the culture (cells µl^−1^) to calculate the total volume (V) of each species in a mixed-algae diet. Tray flow rates (mean flow rate, approx. 480 ± 9 ml^−1^ min^−1^) and food delivery were measured and adjusted daily.

All geoduck survived the exposure periods. Half of the remaining juveniles burrowed in each tray were maintained at the hatchery, positioned in the same replicate trays and stacked for continuous and high flow of ambient seawater (~8-10 L minute^−1^). Stacked trays, commonly used for incubation of finfish, present a promising innovation for geoduck aquaculture; the experiment stack occurred alongside prototype stacked growing trays stocked by Jamestown Point Whitney Shellfish. The juveniles were fed cultured algae *ad libitum* daily for 157 days before shell length and respiration rates were measured.

### 2.2. Respirometry and shell length measurements

Juvenile geoduck were measured on days 2, 5, 8, and 10 of initial exposure, days 0, 2, 4, and 6 (cumulatively as day 24, 26, 28, and 30, respectively) of secondary exposure, and 157 days after the exposure period (cumulatively as day 187) to assess rates of oxygen consumption normalized to shell length. Calibrated optical sensor vials (PreSens, SensorVial SV-PSt5-4ml) were used to measure oxygen consumption in 4 ml vials on a 24-well plate sensor system (Presens SDR SensorDish). Juveniles in each treatment dish were divided into three sensor vials (10 individuals vial^−1^ for exposure periods; 1 individual vial^−1^ at 157-d post-exposure), each filled with 0.2 µm-filtered seawater from corresponding trays. Three blank vials per tray, filled only with 0.2 µm-filtered seawater, were used to account for potential microbial oxygen consumption. Respiratory runs occurred within an incubator at 15°C, with the vials and sensor placed on a rotator for mixing. Each set of measurements lasted ~30 minutes and trials ceased when oxygen concentration declined ~70-80% saturation to avoid hypoxic stress and isolate the effect of *p*CO_2_ treatment on respiration rate. Siphons were observed pre and post-respirometry and were fully extended (~1-2 times shell length). Geoduck were subsequently photographed and shell length (parallel to hinge) was measured using Image J with a size standard (1 mm stage micrometer).

Rates of respiration (oxygen consumption) were calculated from repeated local linear regressions using the R package LoLinR (Olito *et al.*, 2017). An initial criterion of fixed constants (from the LoLin R package) for weighting method (L_%_) and observations (alpha = 0.2) was run individually for each respirometry measurement over the full 30-minute record as a “reference” dataset. These are considered to be the most robust parameters as suggested by the R package authors (Olito *et al.*, 2017). Diagnostic plots (from the LoLin R package) were individually observed and L_%_ and alpha were altered as necessary to best approximate the peak empirical distribution of local linear regressions (see https://doi.org/10.5281/zenodo.3588326 for full details). To determine the optimal set of parameters, respiration data was calculated using three alpha values and data truncations (alpha = 0.2, 0.4, and 0.6; truncation = 10-20 minutes, 10-25 minutes, and no truncation; weighting method = L_%_) and each was compared to the initial reference dataset with two curve fitting steps (local polynomial regressions) to calculate unbiased and reproducible rates of oxygen consumption similar to the reference (10-day exposure, r^2^=0.88; 6-day exposure, r^2^=0.95). Final metabolic rates of juvenile geoduck were corrected for vial volume, rates of oxygen change in the blank vials, and standardized by mean shell length (µg O_2_ hr^−1^ mm^−1^).

### 2.3. Seawater carbonate chemistry

Total alkalinity (TA; µmol kg ^−1^ seawater) water samples were collected from trays once daily during treatment periods, in combination with measurements of pH by handheld probe (Mettler Toledo pH probe; resolution: 1 mV, 0.01 pH; accuracy: ± 1 mV, ± 0.01 pH; Thermo Scientific Orion Star A series A325), salinity (Orion 013010MD Conductivity Cell; range 1 µS/cm −200 mS/cm; accuracy: ± 0.01 psu), and temperature (Fisherbrand Traceable Platinum Ultra-Accurate Digital Thermometer; resolution; 0.001°C; accuracy: ± 0.05 °C). Seawater chemistry was measured for three consecutive days during the 14 days of ambient common garden between initial and secondary treatment periods. Quality control for pH data was assessed daily with Tris standard (Dickson Lab Tris Standard Batch T27) and handheld conductivity probes used for discrete measurements were calibrated every three days. TA was measured using an open cell titration (SOP 3b; Dickson *et al.*, 2007) with certified HCl titrant (∼0.1 mol kg^−1^, ∼0.6 mol kg^−1^ NaCl; Dickson Lab) and TA measurements identified <1% error when compared against certified reference materials (Dickson Lab CO_2_ CRM Batches 137 and 168). Seawater chemistry was completed following Guide to Best Practices (Dickson *et al.*, 2007); daily measurements were used to calculate carbonate chemistry, CO_2_, *p*CO_2_, HCO^3-^, CO_3_, and Ω_aragonite_, using the SEACARB package (Gattuso *et al.*, 2018) in R v3.5.1 (R Core Team, 2018).

### 2.4. Data Analysis

A two-way Analysis of Variance (ANOVA) was used to analyze the effect of time (fixed), *p*CO_2_ treatment (fixed), and time×*p*CO_2_ interaction for respiration and shell length during initial exposure. A t-test was used to test the effect of initial *p*CO_2_ treatment on respiration rate and shell length prior to the secondary exposure (last day of ambient common garden, cumulatively day 24, day 0). For the secondary exposure period, a three-way ANOVA was used to test the effects of time (fixed), initial *p*CO_2_ treatment (fixed), secondary *p*CO_2_ treatment (fixed), and their interactions on respiration rate and shell length. No significant differences in seawater chemistry were detected between trays of the same treatment (pH, *p*CO_2_, TA, salinity, and temperature; https://doi.org/10.5281/zenodo.3588326), thus tray effects were assumed negligible. Significant model effects were followed with pairwise comparisons with a Tukey’s *a posteriori* HSD. We used a two-way ANOVA to analyze the effects of initial (fixed) and secondary (fixed) *p*CO_2_ treatments on respiration and shell length after 157 days in ambient conditions. In all cases, model residuals were tested for normality assumptions with visual inspection of diagnostic plots (residual vs. fitted and normal Q-Q; Kozak and Piepho, 2018) and homogeneity of variance was tested with Levene’s test. Model effects using raw data were robust to transformation(s) that resolved normality assumptions via Shapiro-Wilk test. Statistical tests were completed using R (v3.5.1; R Core Team, 2018) and all code is available (https://doi.org/10.5281/zenodo.3588326).

## 3. Results

### 3.1. Exposure 1

The respiration rate of juvenile clams (4.26 ± 0.85 mm shell length; mean ± SD) prior to exposure was 0.29 ± 0.16 µg O_2_ hr^−1^ mm^−1^ (mean ± SD). Elevated *p*CO_2_ had a significant effect on respiration rate over the initial 10-day exposure (*p*CO_2_ treatment, *F*_1,88_ = 7.512; P < 0.01) with a 25% reduction (averaged across all days) in respiration rate in elevated *p*CO_2_ treatment relative to ambient (Fig. 2A). Juvenile geoduck grew significantly with time under the initial 10-d exposure (time, *F*_3,949_ = 3.392; P = 0.018) with a 3.6% increase in shell length between days 2 and 10 (Fig. 2B), but there was no effect of *p*CO_2_ treatment on shell length (Table 2). Significant differences in respiration rate from the initial *p*CO_2_ treatment were still apparent after 14 days in ambient common garden and before the onset of the secondary exposure (Table 2 and Fig. 3A). In contrast, there was no significant change in shell length due to initial *p*CO_2_ treatment after 14 days in ambient common garden (Table 2).

**Figure 2.**
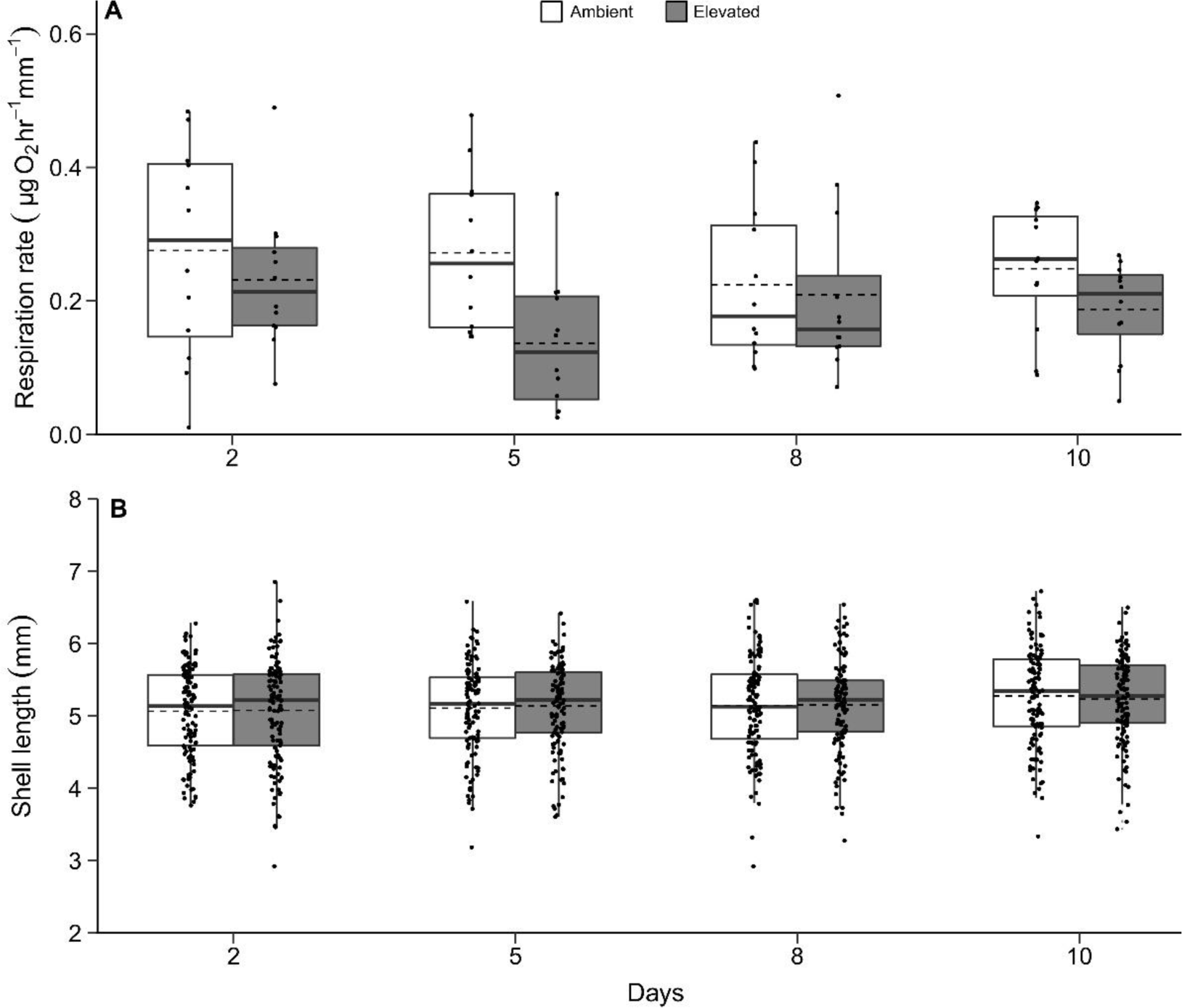
Respiration rates (A) and shell length (B) of juvenile geoduck under the initial 10-day exposure displayed as box whisker plots with mean (dashed line), median (solid line), 25-50% range (box), and interquartile range (whiskers). Small solid circles represent all data points.

**Figure 3.**
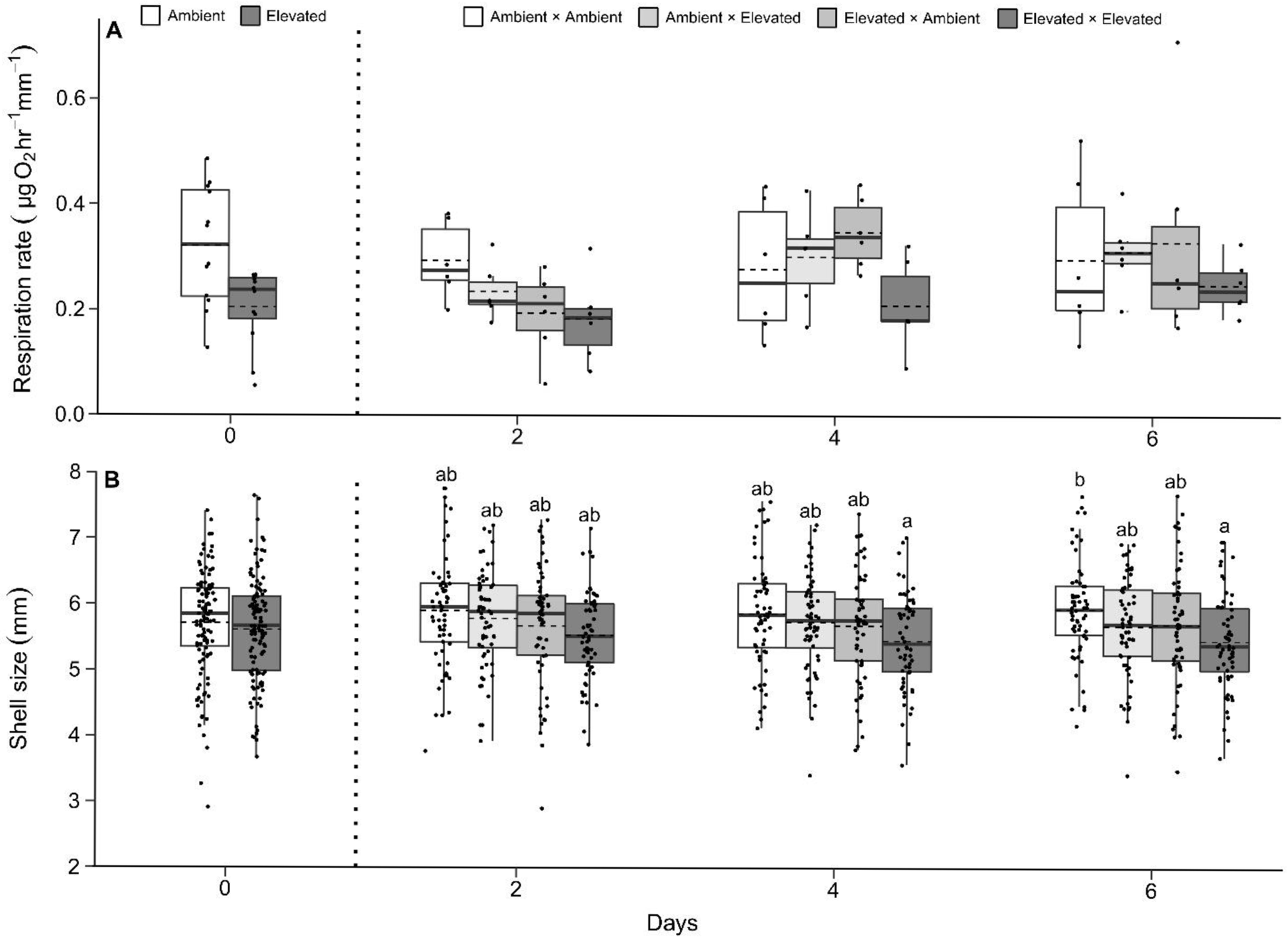
Respiration rates (A) and shell length (B) of juvenile geoduck under the secondary 6-day exposure displayed as box whisker plots with mean (dashed line), median (solid line), 25-50% range (box), and interquartile range (whiskers). Small solid circles represent all data points. Letters display significant post hoc effects and an asterisk shows difference via t-test (P < 0.05).

### 3.2. Exposure 2

There was no interaction between initial and secondary *p*CO_2_ treatments nor between treatments and time on respiration rate or shell length (Table 2). There was a marginal effect of time on respiration rate (Table 2; time, *F*_2,60_ = 3.137; P = 0.0506) with a 31% increase in average respiration rate between days 2 and 6. Initial *p*CO_2_ treatment had a significant effect on shell length, with on average a ~4% reduction in shell size under high *p*CO_2_ relative to ambient initial exposure (Fig. 3B; *p*CO_2_initial_, *F*_1,709_ = 15.821; P < 0.001). This same trend was present under the secondary high *p*CO_2_ exposure, (Fig. 3B; *p*CO_2_secondary_, *F*_1,709_ = 9.917; P = 0.002) with 3.20% smaller shells for individuals exposed to elevated *p*CO_2_ treatments. There were pairwise differences in shell size between animals only exposed to ambient and animals repeatedly exposed to elevated *p*CO_2_ (Fig. 3B; day 6, P = 0.0415; day 6 ambient - day 4 elevated, P = 0.0406).

### 3.3. Common garden after exposure periods

There was no interaction between initial and secondary *p*CO_2_ treatments on respiration rate or shell length (Table 2). The initial exposure period had a significant effect on shell length of juveniles previously exposed to high *p*CO_2_, after 157 days in ambient common garden (Fig. 4A; *p*CO_2_initial_, *F*_1,170_ = 5.228; P = 0.023), where average shell lengths were 5.8% larger in juveniles exposed to initial elevated *p*CO_2_. Secondary 6-day exposure had a significant effect on respiration rates after 157 days in ambient common garden (Fig. 4B; *p*CO_2_seccondary_, *F*_1,31_ =13.008; P = 0.001) with an average of 52.4% greater respiration rates in juveniles secondarily exposed to elevated *p*CO_2_. Visual examination during screening indicated low mortality (1-4 tray^−1^) over the ~5-month grow-out period. Shell lengths of dead animals (as empty shells) were similar to the size of juvenile geoduck during the 30-d exposure period suggesting low mortality occurred at the start of the grow-out period possibly due to handling stress.

**Figure 4.**
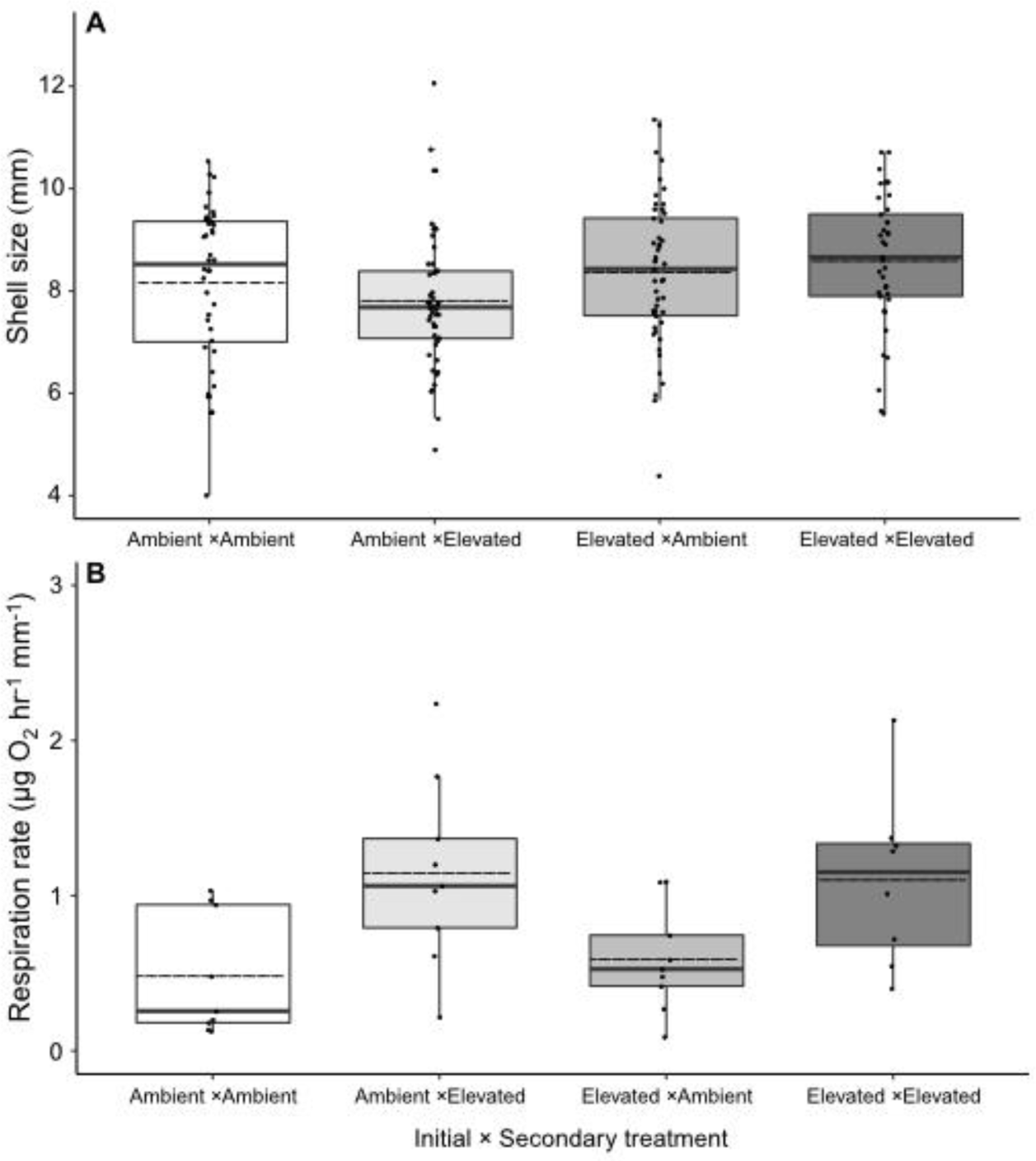
Shell length (A) and metabolic rates (B) of juvenile geoduck after 157 days in ambient common garden conditions post-exposure. Data is displayed as box whisker plots with mean (dashed line), median (solid line), 25-50% range (box), and interquartile range (whiskers). Small solid circles represent all data points.

## 4. Discussion

Metabolic recovery and compensatory shell growth by juvenile *P. generosa* present a novel application of hormetic framework for resilience of a mollusc to acidification. To date, within-generation carry-over effects remain poorly understood for marine molluscs (Ross *et al.*, 2016) with few examples of either positive and negative responses after stress challenges *(Hettinger et al., 2012; Gobler and Talmage, 2013; Putnam et al., 2017).* Further study on conditioning-hormesis in response to *p*CO_2_ stress must address cellular-level energy allocation, in addition to whole organism physiology, to account for essential functions with more holistic implications for stress resilience (Pan *et al.* 2015).

### 4.1. Metabolic depression and compensatory response

Metabolic depression, such that was found under initial exposure of geoduck to elevated *p*CO_2_, has been suggested as an adaptive mechanism to extend survival (Guppy and Withers, 1999). Stress-induced metabolic depression has been documented for a variety of marine invertebrates in response to environmental stress. For example, in the New Zealand geoduck, *Panopea zelandica*, there was a 2-fold reduction in respiration rate under hypoxia (Le *et al.*, 2016). Prior work has shown metabolic reductions up to 60-95% of basal performance at rest for marine molluscs (Guppy and Withers, 1999). Here, depression of oxygen consumption rate by juvenile geoduck to ~25% in comparison with rates under ambient conditions suggests *P. generosa* are relatively tolerant to short-term acidification and may have adaptive physiology to cope with environmental acidification and high *p*CO_2_. Responsiveness to acidification is critical for pH-tolerant taxa to maintain buffering capacity and cope with acidosis (high intracellular *p*CO_2_; (Melzner *et al.*, 2009). However, pH-induced metabolic depression to a similar degree found in this study has caused a permanent decrease in extracellular pH and increase in protein degradation and ammonia excretion in the Mediterranean mussel (*Mytilus galloprovincialis*) *(Michaelidis et al., 2005)*. Conversely, metabolic elevation is relatively common for early-life stage bivalves exposed to low pH and Ω_aragonite_ undersaturation and typically coincides with consequences for performance and survival (Michaelidis *et al.*, 2005; Beniash *et al.*, 2010; Thomsen and Melzner, 2010; Fernández-Reiriz *et al.*, 2011; Waldbusser *et al.*, 2015; Lemasson *et al.*, 2018). Whether depressed or elevated, stress-induced metabolic alterations are known to contribute to biochemical outcomes such as intracellular hypercapnia and hemolymph acidosis (Pörtner *et al.*, 2004; Spicer *et al.*, 2011) and increased ammonia excretion and reduced growth for invertebrate fauna (Michaelidis *et al.*, 2005; Beniash *et al.*, 2010; Lannig *et al.*, 2010; Thomsen and Melzner, 2010; Gazeau *et al.*, 2013). However, *p*CO_2_ did not impair shell growth during the initial period further demonstrative of the pH/hypercapnia-tolerance of *P. generosa*.

Juvenile geoduck repeatedly exposed to elevated *p*CO_2_ showed possible stress “memory” with rebound from metabolic depression under subsequent stress and higher respiration rate and compensatory shell growth after long-term recovery. Metabolic rebound supports a hormetic-like response by *P. generosa* (Calabrese *et al.*, 2007; Costantini, 2014) and prompts further investigation of energy budget, cellular, and -omic measures under repeated reciprocal stress encounters to improve our understanding of the mechanism underpinning hormesis. Use of hormesis to conceptualize carry-over effects of mild stress exposure is largely confined to model insects, plants, and microorganisms (Lee *et al.*, 1987; Calabrese and Blain, 2009; López-Martínez and Hahn, 2012; Visser *et al.*, 2018). For example, Visser et al. (2018) found the Caribbean fruit fly, *Anastrepha suspensa*, exposed to oxidative stress early in life enhanced survivorship and investment in fertility and lipid synthesis under subsequent stress during adulthood. Mechanistic molecular and biochemical assessments under different and repeated stress intensities (i.e. magnitude, duration, and frequency) are planned to determine the threshold between low-dose stimulation and high-dose inhibition from stress-conditioning.

### 4.2. Age and intensity dependence of shell growth

Metabolic recovery was coupled with reduced shell growth under a repeated stress encounter (Fig. 3) and compensatory shell growth after approximately five months in ambient conditions (Fig. 4). This could be explained by several hypotheses such as: carry-over effect from metabolic depression under initial exposure to elevated *p*CO_2_ (Fig. 2A), differing sensitivity to stress intensity (Table 1), and/or age dependence for environmental hardening, or the interaction with increasing temperature through the season (see Supplementary Figure 1.). Bivalves known to exhibit metabolic suppression under acute and long-term acidification are often attributed with increased ammonia excretion rates and decreased ingestion and clearance rates as possible contributors to protein degradation and reduced growth (Michaelidis *et al.*, 2005; Thomsen and Melzner, 2010; Fernández-Reiriz *et al.*, 2011; Navarro *et al.*, 2013). Therefore, decreased shell length under secondary exposure may be a latent effect of metabolic depression during initial exposure. However, shell length was also reduced for clams initially exposed to the elevated treatment in the second exposure period (Table 2, Fig. 3B), indicating potential age-dependence of calcification and bioenergetic effects for juvenile *P. generosa.* This reduction however, could also be explained by the fact the secondary elevated *p*CO_2_ treatment was on average ~0.04 pH units lower than the initial exposure (Table 1), suggesting possible sensitivity to increased stress intensity. It is likely that both temporal dynamics and stress thresholds influence intragenerational carry-over effects and further experimental efforts with repeated reciprocal design are needed.

Respiration rates and shell growth five months post-exposure show a latent enhancement for animals repeatedly stressed or exposed to a stress event earlier in life, emphasizing the importance of the severity, duration, and timing of intragenerational stress-conditioning. These specific findings present a window in their life history where it may be advantageous to condition Pacific geoduck for enhancement of sustainable aquaculture.

### 4.3. Commercial and environmental applications of experimental findings

Our findings infer both positive and negative implications for aquaculture. Although advantageous to elicit carry-over effects exhibited by stress-conditioned animals, results imply greater feed (ingestion rate) to sustain enhanced aerobic metabolism and compensatory shell growth; this can heighten labor and financial costs for industry, likely not incentivized by a marginal 5.8% increase in shell size. However, typical protocols for geoduck aquaculture yield 5-month-old juvenile clams in the hatchery before grown *in-situ* for 5-7 years. Consequently, latency of enhanced performance in this study (~9-month-old juveniles), overlaid with the standard timeline for geoduck industry, does not present additional expenses. Further related tests on stress conditioning and production of resilient strains (i.e. phenotypes and/or epigenotypes) must account for distinct life-stages and species-specific attributes in aquaculture practice.

Shellfish farming has adapted in recent years to implement “climate-proofing” technology to maintain production and combat both coastal and climate-related stressors (e.g. ocean acidification, sea-level rise, coastal development; Allison et al. 2011). For example, chemical buffering systems (e.g. mixing sodium bicarbonate) are increasingly common in shellfish industry to elevate aragonite saturation levels and reduce deleterious effects of ocean acidification; hatcheries report increases in productivity by 30-50%, offsetting the cost to maintain optimal carbonate chemistry year-round (Barton et al. 2015). Although buffering systems are advantageous to yield juvenile ‘seed’, alleviation of aragonite undersaturation in the short-term may leave juveniles and adults unprepared to cope with the heterogeneity of environmental chemistry during long growing periods *in-situ*. As conditions in coastal bays report deteriorating water quality (Feely et al. 2008; Wallace et al. 2014; Cloern 2001; Melzner et al. 2013), acclimatization and selective breeding posit alternate and more robust solutions to generate stress-resilience (Barton et al. 2015). Implementation and tests of effectiveness of stress conditioning remain uncommon for scientists and aquaculture; our novel findings collected in a hatchery setting provide incentive to fine-tune stress exposures and build a mechanistic understanding of physiological, cellular, and molecular responses. Critical questions to test the practical application of stress conditioning are: (1) what are the effects of repeated stress exposures on energy budget? (2) what life-stages and/or *p*CO_2_ stress intensity (i.e. magnitude and duration) optimizes establishment of resilient phenotypes and genotypes during hatchery-rearing? (3) does stress history under elevated *p*CO_2_ affect the stability and longevity of carry-over effects later in life? Answers to these challenges will result in effective implementation of conditioning to both reduce pressure on wild stocks and sustain food security under environmental change.

Although this study was primarily focused on production enhancement in a hatchery setting, effects on shell growth and metabolism have important applications to natural systems. Seawater carbonate chemistry targeted for stress treatments was more severe than levels commonly used in experimental research (Gazeau *et al.*, 2010; Navarro *et al.*, 2013; Diaz *et al.*, 2018), but relevant to summer subsurface conditions within the natural range of *P. generosa* (pH 7.4 and Ω_aragonite_ 0.4 in Hood Canal, WA; Feely *et al.*, 2010). Thus, survival, metabolic recovery, and compensatory growth in *P. generosa* in this study demonstrates a resilience to short-term acidification in the water column. Enhanced growth rates during juvenile development can present benefits for burrowing behavior (Green *et al.*, 2009; Clements *et al.*, 2016; Meseck *et al.*, 2018) and survival due to decreased risk of predation and susceptibility to environmental stress (Przeslawski and Webb, 2009; Johnson and Smee, 2012). Specific to juvenile *P. generosa*, time to metamorphosis (to dissoconch), pre-burrowing time (time elapsed to anchor into substrate and obtain upright position), and burrowing depth are directly related to growth and survival (Goodwin and Pease, 1989; Tapia-Morales *et al.*, 2015). Thus, stress conditioning under CO_2_-enrichment and low pH may enhance survivorship of juvenile geoduck in natural systems. Water column carbonate chemistry may be critical for sustainable production of infaunal clams, such as *P. generosa*, that are out-planted for several years *in-situ* on mudflats known to exhibit dynamic abiotic gradients (Green *et al.*, 1993; Burdige *et al.*, 2008) adjacent to seasonally acidified and undersaturated water bodies (Feely *et al.*, 2010; Reum *et al.*, 2014).

## Conclusion

Data in this present study provides evidence of capacity to cope with short-term acidification for an understudied infaunal clam of high economic importance. Survival of all individuals over the 30-d experiment demonstrates the resilience of this species to low pH and reduced carbonate saturation. Juvenile geoduck exposed to low pH for 10 days recovered from metabolic depression under subsequent stress exposure and conditioned animals showed a significant increase in both shell length and metabolic rate compared to controls after five months under ambient conditions, suggesting stress “memory” and compensatory growth as possible indicators of enhanced performance from intragenerational stress-conditioning. Our focus on industry enhancement must expand to test developmental morphology, physiology, and genetic and non-genetic markers over larval and juvenile stages in a multi-generational experiment to generate a more holistic assessment of stress hardening and the effects of exposure on cellular stress response (Costantini *et al.*, 2010; Foo and Byrne, 2016; Eirin-Lopez and Putnam, 2018) for advancement of sustainable aquaculture (Branch *et al.*, 2013). Advancements in genome sequencing will facilitate further research to synthesize -omic profiling (i.e global DNA methylation and differential expression) with physiological responses throughout reproductive and offspring development under environmental stress (Gavery and Roberts, 2014; Li *et al.*, 2019) to determine if these mechanisms are transferable among species. Stress conditioning within a generation at critical life stages may yield beneficial responses for food production and provide a baseline for other long-lived burrowing bivalves of ecological and economic importance.

## Supplementary Material

Supplementary Figure 1. Continuous temp and pH data from APEX system

## Acknowledgements

We thank the Jamestown S’Klallam Tribe and Kurt Grinnell for providing the animals and facilities for this research. We also thank the management staff and technicians at the Jamestown Point Whitney Shellfish Hatchery, Matt Henderson, Josh Valley, and Clara Duncan, for their assistance, and Maddie Sherman, Emma Strand, and Kaitlyn Mitchell for analytical support.

---

**Supplementary Figure 1.**
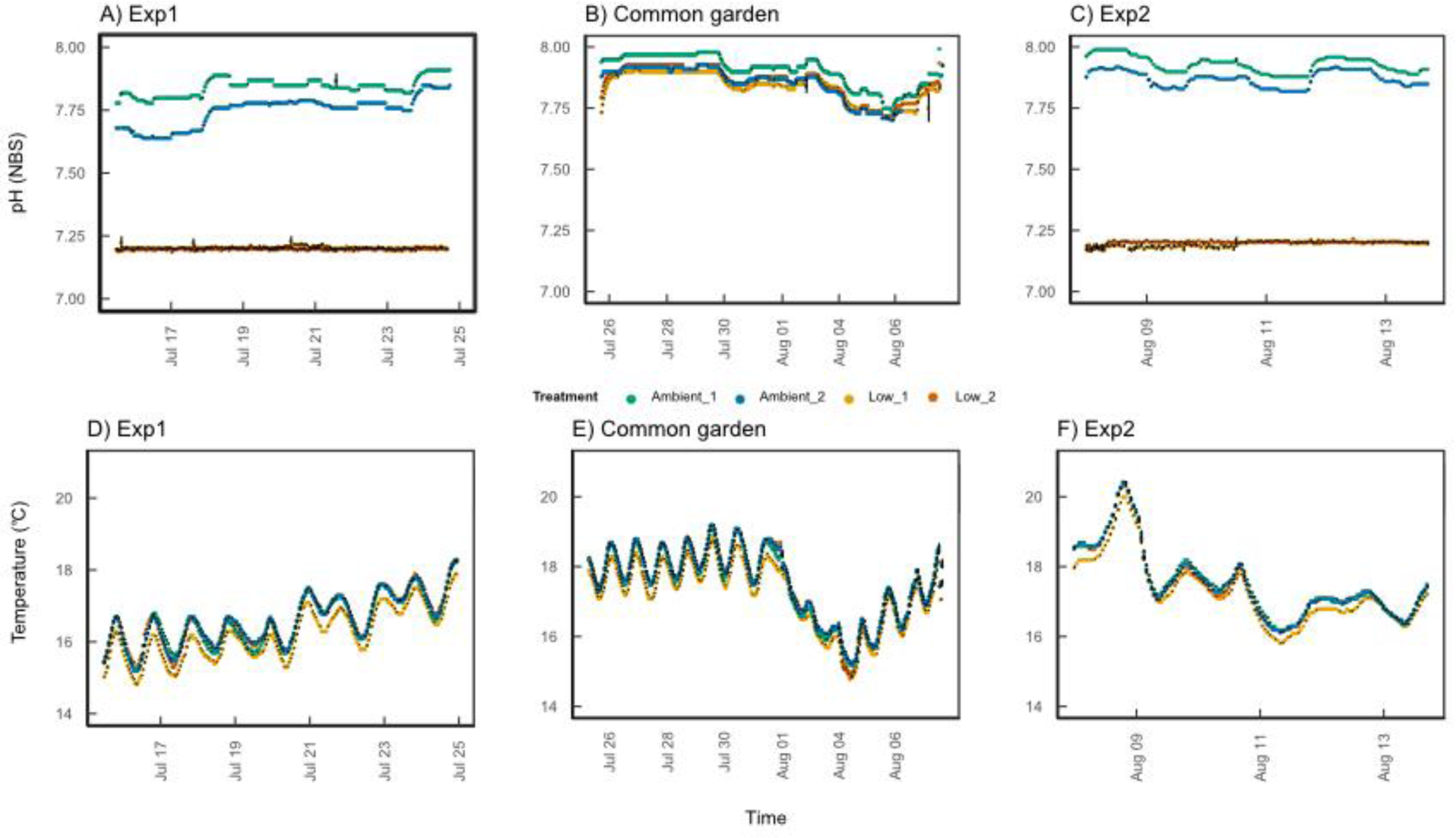
Hourly averages (mean ± SEM) of temperature (°C) and pH (NBS scale) data from the Neptune Apex system. Data were collected every 10 minutes.

## References

Allison EH, Badjeck M-C, Meinhold K (2011) The Implications of Global Climate Change for Molluscan Aquaculture. Shellfish Aquaculture and the Environment.

Barton A, Hales B, Waldbusser GG, Langdon C, Feely RA (2012) The Pacific oyster, Crassostrea gigas, shows negative correlation to naturally elevated carbon dioxide levels: Implications for near-term ocean acidification effects. Limnol Oceanogr 57: 698–710.

Barton A, Hatchery WCS, Waldbusser G, Feely R, Weisberg S, Newton J, Hales B, Cudd S, Eudeline B, Langdon C, et al. (2015) Impacts of coastal acidification on the Pacific Northwest shellfish industry and adaptation strategies implemented in response. Oceanography 25: 146–159.

Beniash E, Ivanina A, Lieb NS, Kurochkin I, Sokolova IM (2010) Elevated level of carbon dioxide affects metabolism and shell formation in oysters Crassostrea virginica (Gmelin). Mar Ecol Prog Ser 419: 95–108.

Branch TA, DeJoseph BM, Ray LJ, Wagner CA (2013) Impacts of ocean acidification on marine seafood. Trends Ecol Evol 28: 178–186.

Burdige DJ, Zimmerman RC, Hu X (2008) Rates of carbonate dissolution in permeable sediments estimated from pore-water profiles: The role of sea grasses. Limnol Oceanogr 53: 549–565.

Cai W-J, Hu X, Huang W-J, Murrell MC, Lehrter JC, Lohrenz SE, Chou W-C, Zhai W, Hollibaugh JT, Wang Y, et al. (2011) Acidification of subsurface coastal waters enhanced by eutrophication. Nat Geosci 4: 766–770.

Calabrese EJ (2008) Hormesis: why it is important to toxicology and toxicologists. Environ Toxicol Chem 27: 1451–1474.

Calabrese EJ, Bachmann KA, Bailer AJ, Bolger PM, Borak J, Cai L, Cedergreen N, Cherian MG, Chiueh CC, Clarkson TW, et al. (2007) Biological stress response terminology: Integrating the concepts of adaptive response and preconditioning stress within a hormetic dose-response framework. Toxicol Appl Pharmacol 222: 122–128.

Calabrese EJ, Blain RB (2009) Hormesis and plant biology. Environmental Pollution.

Calabrese EJ, Mattson MP (2011) Hormesis provides a generalized quantitative estimate of biological plasticity. J Cell Commun Signal 5: 25–38.

Campbell A, Harbo RM, Hand CM (1998) Harvesting and distribution of Pacific geoduck clams, Panopea abrupta, in Brtitish Columbia. Can Spec Publ Fish Aquat Sci/Publ Spec Can Sci Halieut. Aquat 125: 349–358.

Cao R, Liu Y, Wang Q, Yang D, Liu H, Ran W, Qu Y, Zhao J (2018) Seawater acidification reduced the resistance of Crassostrea gigas to Vibrio splendidus challenge: an energy metabolism perspective. Frontiers in Physiology.

Clements JC, Woodard KD, Hunt HL (2016) Porewater acidification alters the burrowing behavior and post-settlement dispersal of juvenile soft-shell clams (Mya arenaria). J Exp Mar Bio Ecol 477: 103–111.

Cloern JE (2001) Our evolving conceptual model of the coastal eutrophication problem. Marine Ecology Progress Series.

Cole VJ, Parker LM, O’Connor SJ, O’Connor WA, Scanes E, Byrne M, Ross PM (2016) Effects of multiple climate change stressors: ocean acidification interacts with warming, hyposalinity, and low food supply on the larvae of the brooding flat oyster Ostrea angasi. Mar Biol 163. doi:10.1007/s00227-016-2880-4

Costantini D (2014) Does hormesis foster organism resistance to extreme events? Frontiers in Ecology and the Environment.

Costantini D (2019) Hormesis promotes evolutionary change. Dose Response 17: 1–4.

Costantini D, Metcalfe NB, Monaghan P (2010) Ecological processes in a hormetic framework. Ecol Lett 13: 1435–1447.

Costantini D, Monaghan P, Metcalfe NB (2012) Early life experience primes resistance to oxidative stress. J Exp Biol 215: 2820–2826.

Cubillo AM, Ferreira JG, Pearce CM, Marshall R, Cheney D, Hudson B (2018) Ecosystem services of geoduck farming in South Puget Sound, USA: a modeling analysis. Aquac Int 26: 1427–1443.

Dethier M (2006) Native Shellfish in Nearshore Ecosystems of Puget Sound. Puget Sound Nearshore Partnership Report No. 2006-04. Seattle District, U.S. Army Corps of Engineers, Seattle, Washington.

Diaz RJ, Rosenberg R (2001) Overview of anthropogenically-induced hypoxic effects on marine benthic fauna. Coastal and Estuarine Studies.

Diaz R, Lardies MA, Tapia FJ, Tarifeño E, Vargas CA (2018) Transgenerational effects of pCO_2_-driven ocean acidification on adult mussels Mytilus chilensis modulate physiological response to multiple stressors in larvae. Front Physiol 9. doi:10.3389/fphys.2018.01349

Dickson AG, Sabine CL, Christian JR, eds. (2007) Guide to Best Practices for Ocean CO_2_ Measurements. PICES.

Dupont S, Dorey N, Stumpp M, Melzner F, Thorndyke M (2012) Long-term and trans-life-cycle effects of exposure to ocean acidification in the green sea urchin Strongylocentrotus droebachiensis. Mar Biol 160: 1835–1843.

Dupont S, Thorndyke MC (2009) Impact of CO_2_-driven ocean acidification on invertebrates early life-history – What we know, what we need to know and what we can do. Biogeosciences Discussions.

Eirin-Lopez JM, Putnam HM (2018) Marine environmental epigenetics. Ann Rev Mar Sci. doi:10.1146/annurev-marine-010318-095114

Elston RA, Hasegawa H, Humphrey KL, Polyak IK, Häse CC (2008) Re-emergence of Vibrio tubiashii in bivalve shellfish aquaculture: severity, environmental drivers, geographic extent and management. Dis Aquat Organ 82: 119–134.

FAO (2018) The State of World Fisheries and Aquaculture 2018 - Meeting the Sustainable Development Goals.

Feely RA, Alin SR, Newton J, Sabine CL, Warner M, Devol A, Krembs C, Maloy C (2010) The combined effects of ocean acidification, mixing, and respiration on pH and carbonate saturation in an urbanized estuary. Estuar Coast Shelf Sci 88: 442–449.

Fernández-Reiriz MJ, Range P, Álvarez-Salgado XA, Labarta U (2011) Physiological energetics of juvenile clams Ruditapes decussatus in a high CO_2_ coastal ocean. Mar Ecol Prog Ser 433: 97–105.

Foo SA, Byrne M (2016) Acclimatization and adaptive capacity of marine species in a changing ocean. In: Advances in Marine Biology. pp 69–116.

Froehlich HE, Gentry RR, Halpern BS (2018) Global change in marine aquaculture production potential under climate change. Nat Ecol Evol 2: 1745–1750.

Gracía E, Clemente S, Hernández JC (2018) Effects of natural current pH variability on the sea urchin Paracentrotus lividus larvae development and settlement. Mar Environ Res 139: 11–18.

Gtatuso JP, Epitalon JM, Lavine H (2018) Seacarb: Seawater Carbonate Chemistry.

Gvaery MR, Roberts SB (2014) A context dependent role for DNA methylation in bivalves. Brief Funct Genomics 13: 217–222.

Gzaeau F, Gattuso JP, Dawber C, Pronker AE, Peene F, Peene J, Heip CHR, Middelburg JJ (2010) Effect of ocean acidification on the early life stages of the blue mussel (Mytilus edulis). Biogeosci Discuss 7: 2927–2947.

Gzaeau F, Parker LM, Comeau S, Gattuso J-P, O’Connor WA, Martin S, Pörtner H-O, Ross PM (2013) Impacts of ocean acidification on marine shelled molluscs. Mar Biol 160: 2207–2245.

Gimenez I, Waldbusser GG, Hales B (2018) Ocean acidification stress index for shellfish (OASIS): Linking Pacific oyster larval survival and exposure to variable carbonate chemistry regimes. Elem Sci Anth 6: 51.

Gobler CJ, Talmage SC (2013) Short- and long-term consequences of larval stage exposure to constantly and ephemerally elevated carbon dioxide for marine bivalve populations. Biogeosciences.

Goncalves P, Anderson K, Raftos DA, Thompson EL (2018) The capacity of oysters to regulate energy metabolism-related processes may be key to their resilience against ocean acidification. Aquac Res 49: 2059–2071.

Goodwin L, Pease B (1989) Species Profiles: Life Histories and Environmental Requirements of Coastal Fish and Invertebrates (Pacific Northwest): Pacific Geoduck Clam. US Fish and Wildlife.

Green MA, Aller RC, Aller JY (1993) Carbonate dissolution and temporal abundances of Foraminifera in Long Island Sound sediments. Limnol Oceanogr 38: 331–345.

Green MA, Waldbusser GG, Reilly SL, Emerson K, O’Donnell S (2009) Death by dissolution: Sediment saturation state as a mortality factor for juvenile bivalves. Limnol Oceanogr 54: 1037–1047.

Gunderson AR, Armstrong EJ, Stillman JH (2016) Multiple stressors in a changing world: the need for an improved perspective on physiological responses to the dynamic marine environment. Ann Rev Mar Sci 8: 357–378.

Guppy M, Withers P (1999) Metabolic depression in animals: physiological perspectives and biochemical generalizations. Biol Rev Camb Philos Soc 74: 1–40.

Gurr SJ, Rollando C, Chan LL, Vadopalas B, Putnam HM, Roberts SB (2018) Alternative Image-Based Technique for Phytoplankton Cell Counts in Shellfish Aquaculture (No. 1001481). Nexcelom.

Hettinger A, Sanford E, Hill TM, Russell AD, Sato KNS, Hoey J, Forsch M, Page HN, Gaylord B (2012) Persistent carry-over effects of planktonic exposure to ocean acidification in the Olympia oyster. Ecology 93: 2758–2768.

Johnson KD, Smee DL (2012) Size matters for risk assessment and resource allocation in bivalves. Marine Ecology Progress Series.

Kapsenberg L, Miglioli A, Bitter MC, Tambutté E, Dumollard R, Gattuso JP (2018) Ocean pH fluctuations affect mussel larvae at key developmental transitions. Proceedings of the Royal Society B: Biological Sciences 285: 20182381.

Kozak M, Piepho HP (2018) What’s normal anyway? Residual plots are more telling than significance tests when checking ANOVA assumptions. Journal of Agronomy and Crop Science.

Kroeker KJ, Kordas RL, Crim RN, Singh GG (2010) Meta-analysis reveals negative yet variable effects of ocean acidification on marine organisms. Ecol Lett 13: 1419–1434.

Lannig G, Eilers S, Pörtner HO, Sokolova IM, Bock C (2010) Impact of ocean acidification on energy metabolism of oyster, Crassostrea gigas—changes in metabolic pathways and thermal response. Marine Drugs.

Le DV, Alfaro AC, Ragg NLC, Hilton Z, King N (2016) Aerobic scope and oxygen regulation of New Zealand geoduck (Panopea zelandica) in response to progressive hypoxia. Aquaculture 463: 28–36.

Lee RE Jr, Chen CP, Denlinger DL (1987) A rapid cold-hardening process in insects. Science 238: 1415–1417.

Lemasson AJ, Hall-Spencer JM, Fletcher S, Provstgaard-Morys S, Knights AM (2018) Indications of future performance of native and non-native adult oysters under acidification and warming. Mar Environ Res 142: 178–189.

Li Y, Zhang L, Li Y, Li W, Guo Z, Li R, Hu X, Bao Z, Wang S (2019) Dynamics of DNA methylation and DNMT expression during gametogenesis and early development of scallop Patinopecten yessoensis. Mar Biotechnol. doi:10.1007/s10126-018-09871-w

Lohbeck KT, Riebesell U, Reusch TBH (2012) Adaptive evolution of a key phytoplankton species to ocean acidification. Nat Geosci 5: 346–351.

López-Martínez G, Hahn DA (2012) Short-term anoxic conditioning hormesis boosts antioxidant defenses, lowers oxidative damage following irradiation and enhances male sexual performance in the Caribbean fruit fly, Anastrepha suspensa. J Exp Biol 215: 2150–2161.

Mackenzie CL, Ormondroyd GA, Curling SF, Ball RJ, Whiteley NM, Malham SK (2014) Ocean warming, more than acidification, reduces shell strength in a commercial shellfish species during food limitation. PLoS ONE.

Mangi SC, Lee J, Pinnegar JK, Law RJ, Tyllianakis E, Birchenough SNR (2018) The economic impacts of ocean acidification on shellfish fisheries and aquaculture in the United Kingdom. Environ Sci Policy 86: 95–105.

Melzner F, Gutowska MA, Langenbuch M, Dupont S, Lucassen M, Thorndyke MC, Bleich M, Pörtner HO (2009) Physiological basis for high CO_2_ tolerance in marine ectothermic animals: pre-adaptation through lifestyle and ontogeny? Biogeosci Discuss 6: 4693–4738.

Meseck SL, Mercaldo-Allen R, Kuropat C, Clark P, Goldberg R (2018) Variability in sediment-water carbonate chemistry and bivalve abundance after bivalve settlement in Long Island Sound, Milford, Connecticut. Mar Pollut Bull 135: 165–175.

Michaelidis B, Ouzounis C, Paleras A, Pörtner HO (2005) Effects of long-term moderate hypercapnia on acid-base balance and growth rate in marine mussels Mytilus galloprovincialis. Mar Ecol Prog Ser 293: 109–118.

Navarro JM, Torres R, Acuña K, Duarte C, Manriquez PH, Lardies M, Lagos NA, Vargas C, Aguilera V (2013) Impact of medium-term exposure to elevated pCO_2_ levels on the physiological energetics of the mussel Mytilus chilensis. Chemosphere 90: 1242–1248.

Olito C, White CR, Marshall DJ, Barneche DR (2017) Estimating monotonic rates from biological data using local linear regression. J Exp Biol 220: 759–764.

Orensanz JM (lobo), Hand CM, Parma AM, Valero J, Hilborn R (2004) Precaution in the harvest of Methuselah’s clams the difficulty of getting timely feedback from slow-paced dynamics. Can J Fish Aquat Sci 61: 1355–1372.

Pan T-CF, Applebaum SL, Manahan DT (2015) Experimental ocean acidification alters the allocation of metabolic energy. Proc Natl Acad Sci USA 112: 4696–4701.

Parker LM, O’Connor WA, Byrne M, Coleman RA, Virtue P, Dove M, Gibbs M, Spohr L, Scanes E, Ross PM (2017) Adult exposure to ocean acidification is maladaptive for larvae of the Sydney rock oyster in the presence of multiple stressors. Biol Lett 13. doi:10.1098/rsbl.2016.0798

Parker LM, O’Connor WA, Raftos DA, Pörtner H-O, Ross PM (2015) Persistence of positive carryover effects in the oyster, Saccostrea glomerata, following transgenerational exposure to ocean acidification. PLoS One 10: e0132276.

Parker LM, Ross PM, O’Connor WA, Borysko L, Raftos DA, Pörtner H-O (2011) Adult exposure influences offspring response to ocean acidification in oysters. Glob Chang Biol 18: 82–92.

Portner HO, Farrell AP (2008) Physiology and climate change. Science 322: 690–692.

Pörtner HO, Langenbuch M, Reipschläger A (2004) Biological impact of elevated ocean CO_2_ concentrations: lessons from animal physiology and earth history. J Oceanogr 60: 705–718.

Prado S, Romalde JL, Montes J, Barja JL (2005) Pathogenic bacteria isolated from disease outbreaks in shellfish hatcheries. First description of Vibrio neptunius as an oyster pathogen. Dis Aquat Organ 67: 209–215.

Przeslawski R, Webb AR (2009) Natural variation in larval size and developmental rate of the northern quahog Mercenaria mercenaria and associated effects on larval and juvenile fitness. Journal of Shellfish Research.

Putnam HM, Davidson JM, Gates RD (2016) Ocean acidification influences host DNA methylation and phenotypic plasticity in environmentally susceptible corals. Evol Appl 9: 1165–1178.

Putnam HM, Gates RD (2015) Preconditioning in the reef-building coral Pocillopora damicornis and the potential for trans-generational acclimatization in coral larvae under future climate change conditions. J Exp Biol 218: 2365–2372.

Putnam HM, Ritson-Williams R, Cruz JA, Davidson JM, Gates RD (2018) Nurtured by nature: considering the role of environmental and parental legacies in coral ecological performance. bioRxiv. doi:10.1101/317453

Putnam H, Roberts S, Spencer LH (2017) Capacity or adaptation and acclimatization to ocean acidification in geoduck through epigenetic mechanisms.

R Core Team (2018) A Language and Environment for Statistical Computing.

Reum JCP, Alin SR, Feely RA, Newton J, Warner M, McElhany P (2014) Seasonal carbonate chemistry covariation with temperature, oxygen, and salinity in a fjord estuary: implications for the design of ocean acidification experiments. PLoS One 9: e89619.

Ries JB, Cohen AL, McCorkle DC (2009) Marine calcifiers exhibit mixed responses to CO_2_-induced ocean acidification. Geology 37: 1131–1134.

Rojas R, Miranda CD, Opazo R, Romero J (2015) Characterization and pathogenicity of Vibrio splendidus strains associated with massive mortalities of commercial hatchery-reared larvae of scallop Argopecten purpuratus (Lamarck, 1819). J Invertebr Pathol 124: 61–69.

Ross PM, Parker L, Byrne M (2016) Transgenerational responses of molluscs and echinoderms to changing ocean conditions. ICES Journal of Marine Science: Journal du Conseil.

Sanders MB, Bean TP, Hutchinson TH, Le Quesne WJF (2013) Juvenile king scallop, Pecten maximus, is potentially tolerant to low levels of ocean acidification when food is unrestricted. PLoS One 8: e74118.

Scanes E, Parker LM, O’Connor WA, Stapp LS, Ross PM (2017) Intertidal oysters reach their physiological limit in a future high-CO_2_ world. J Exp Biol 220: 765–774.

Shamshak GL, King JR (2015) From cannery to culinary luxury: the evolution of the global geoduck market. Mar Policy 55: 81–89.

Shirayama Y (2005) Effect of increased atmospheric CO_2_ on shallow water marine benthos. J Geophys Res 110. doi:10.1029/2004jc002618

Shumway SE, Davis C, Downey R, Karney R, Kraeuter J, Parsons J, Rheault R, Wikfors G (2003) Shellfish aquaculture–in praise of sustainable economies and environments. World Aquacult 34: 8–10.

Spencer LH, Horwith M, Lowe AT, Venkataraman YR, Timmins-Schiffman E, Nunn BL, Roberts SB (2018) Pacific geoduck (Panopea generosa) resilience to natural pH variation.

Spicer JI, Widdicombe S, Needham HR, Berge JA (2011) Impact of CO_2_-acidified seawater on the extracellular acid–base balance of the northern sea urchin Strongylocentrotus dröebachiensis. Journal of Experimental Marine Biology and Ecology.

Stevens AM, Gobler CJ (2018) Interactive effects of acidification, hypoxia, and thermal stress on growth, respiration, and survival of four North Atlantic bivalves. Mar Ecol Prog Ser 604: 143–161.

Suckling CC, Clark MS, Richard J, Morley SA, Thorne MAS, Harper EM, Peck LS (2015) Adult acclimation to combined temperature and pH stressors significantly enhances reproductive outcomes compared to short-term exposures. J Anim Ecol 84: 773–784.

Talmage SC, Gobler CJ (2010) Effects of past, present, and future ocean carbon dioxide concentrations on the growth and survival of larval shellfish. Proc Natl Acad Sci U S A 107: 17246–17251.

Tapia-Morales S, García-Esquivel Z, Vadopalas B, Davis J (2015) Growth and burrowing rates of juvenile geoducks Panopea generosa and Panopea globosa under laboratory conditions. Journal of Shellfish Research.

Thomsen J, Casties I, Pansch C, Körtzinger A, Melzner F (2013) Food availability outweighs ocean acidification effects in juvenile Mytilus edulis: laboratory and field experiments. Glob Chang Biol 19: 1017–1027.

Thomsen J, Melzner F (2010) Moderate seawater acidification does not elicit long-term metabolic depression in the blue mussel Mytilus edulis. Mar Biol 157: 2667–2676.

Thomsen J, Stapp LS, Haynert K, Schade H, Danelli M, Lannig G, Mathias Wegner K, Melzner F (2017) Naturally acidified habitat selects for ocean acidification–tolerant mussels. Science Advances 3: e1602411.

Utting SD, Millican PF (1997) Techniques for the hatchery conditioning of bivalve broodstocks and the subsequent effect on egg quality and larval viability. Aquaculture 155: 45–54.

Visser B, Williams CM, Hahn DA, Short CA, López-Martínez G (2018) Hormetic benefits of prior anoxia exposure in buffering anoxia stress in a soil-pupating insect. The Journal of Experimental Biology.

Waldbusser GG, Hales B, Langdon CJ, Haley BA, Schrader P, Brunner EL, Gray MW, Miller CA, Gimenez I, Hutchinson G (2015) Ocean acidification has multiple modes of action on bivalve larvae. PLoS One 10: e0128376.

Waldbusser GG, Voigt EP, Bergschneider H, Green MA, Newell RIE (2010) Biocalcification in the eastern oyster (Crassostrea virginica) in relation to long-term trends in Chesapeake Bay pH. Estuaries Coasts 34: 221–231.

Wallace RB, Baumann H, Grear JS, Aller RC, Gobler CJ (2014) Coastal ocean acidification: The other eutrophication problem. Estuarine, Coastal and Shelf Science.

Washington Sea Grant (2015) Shellfish Aquaculture in Washington State. Final report to the Washington State Legislature. Washington Sea Grant.

White MM, McCorkle DC, Mullineaux LS, Cohen AL (2013) Early exposure of bay scallops (Argopecten irradians) to high CO_2_ causes a decrease in larval shell growth. PLoS One 8: e61065.

Zhang Z, Hand C (2006) Recruitment patterns and precautionary exploitation rates for geoduck (Panopea abrupta) populations in British Columbia. J Shellfish Res 25: 445–453.

Zhao L, Schöne BR, Mertz-Kraus R, Yang F (2017) Sodium provides unique insights into transgenerational effects of ocean acidification on bivalve shell formation. Sci Total Environ 577: 360–366.

