## Supplementary figures and images for "Metabolic Recovery and Compensatory Shell Growth of Juvenile Pacific Geoduck *Panopea Generosa* Following Short-Term Exposure to Acidified Seawater"

### Supplementary Figure 1

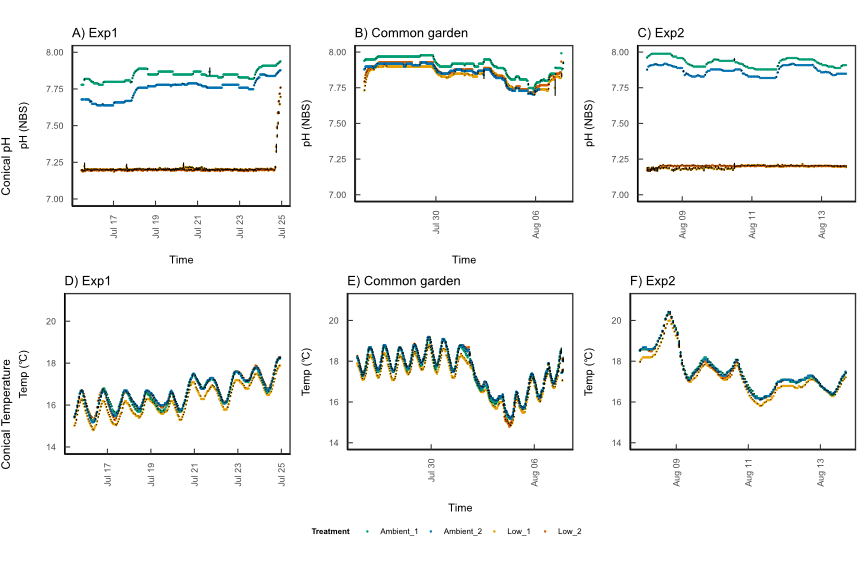
